# Decline in constitutive proliferative activity in the zebrafish retina with ageing

**DOI:** 10.1101/2021.06.16.448637

**Authors:** Ismael Hernández-Núñez, Ana Quelle-Regaldie, Laura Sánchez, Fátima Adrio, Eva Candal, Antón Barreiro-Iglesias

**Author notes:** **Correspondence:** Dr. Antón Barreiro-Iglesias. Equal contributors.

## Abstract

It is largely assumed that the teleost retina shows continuous and active proliferative and neurogenic activity throughout life. But when deepening in the teleost literature one finds that assumptions about a highly active and continuous proliferation in the adult retina are based on studies in which proliferation was not quantified in a comparative way at the different life stages or was mainly studied in juveniles/young adults. Here, we performed a systematic and comparative study of the constitutive proliferative activity of the retina from early developing (2 days post-fertilization) to aged (up to 3-4 years post-fertilization) zebrafish. Mitotic activity and cell cycle progression were analyzed by using immunofluorescence against pH3 and PCNA, respectively. We observed a decline in cell proliferation in the retina with ageing, even despite the occurrence of a wave of secondary proliferation during sexual maturation. During this wave of secondary proliferation the distribution of proliferating and mitotic cells changes from the inner to the outer nuclear layer in the central retina. Importantly, in aged zebrafish there is a virtual disappearance of mitotic activity. Our results showing a decline in proliferative activity of the zebrafish retina with ageing are of crucial importance since it is generally assumed that the fish retina has continuous proliferative activity throughout life.

## 1. Introduction

Neurogenesis is the process by which neural progenitor cells give rise to mature neurons and glial cells. Early in development, the central nervous system (CNS) is formed from a highly active neurogenic neuroepithelium. As development progresses, proliferative and neurogenic activities are gradually lost in most CNS regions, and in postnatal life, neurogenic activity is restricted to specific regions called neurogenic niches (Doetsch, 2003; Álvarez-Buylla and Lim, 2004). Moreover, the presence of postnatal neurogenic activity in the CNS was also progressively lost during vertebrate evolution (reviewed in Ferretti, 2011; Zupanc and Sîrbulescu, 2011; Grandel and Brand, 2013; Than-Trong and Bally-Cuif, 2015; Alunni and Bally-Cuif, 2016; Zupanc, 2021). Accordingly, different vertebrate species show different postnatal/adult proliferative and neurogenic rates and different numbers of neurogenic niches in the CNS, which are more abundant in teleost fishes (reviewed in Ferretti, 2011; Zupanc and Sîrbulescu, 2011; Grandel and Brand, 2013; Than-Trong and Bally-Cuif, 2015; Alunni and Bally-Cuif, 2016; Miles and Tropepe, 2021; Zupanc, 2021). Some postnatal constitutive and/or inducible neurogenic niches are found in the retina of vertebrates. These include the ciliary marginal zone (CMZ), which is a circumferential ring of cells located in the peripheral retina (Harris and Perron, 1998; Raymond et al., 2006; Fisher et al., 2013; Marcucci et al., 2016, Bélanger et al., 2017); the Müller glial cells of the inner nuclear layer (INL) of the central retina (Fausett and Goldman, 2006; Raymond et al., 2006; Bernardos et al., 2007; Nagashima et al., 2013); the retinal pigment epithelium (RPE; Okada, 1980; Engelhardt et al., 2005; Ma et al., 2009); a pseudostratified region at the junction between the retina and the ciliary body (Eymann et al., 2019); and the pigmented and non-pigmented epithelium of the ciliary body (Tropepe et al., 2000; Fischer and Reh, 2001, 2003; Das et al., 2005, 2006). The proliferative and neurogenic capacity of each of these retinal neurogenic niches varies in different vertebrate species (reviewed in Reh and Fischer, 2001; Amato et al., 2004; Moshiri et al., 2004). In fishes, all retinal cell types, except rod photoreceptors, are generated within the CMZ and incorporated to the peripheral retina (so that older cells remain in the central retina and new cells become located successively in more peripheral positions). Instead, rod photoreceptors are continuously generated from Müller glia in the central retina.

Based on studies in teleost species (see references in Table S1), it is largely assumed that the retina of fishes, in contrast to mammals, has continuous proliferative activity throughout life and that this (together with tissue stretching) is partially responsible for continuous eye growth, even during adulthood. This idea emerges in relevant articles on this topic during the last decades: “Fish retinas differ fundamentally from those of other vertebrates because they continue to grow throughout the life of the animal, both by adding new neurons and by stretching existing retinal tissue.” (Fernald, 1991); “In fish and amphibia, retinal stem cells located in the periphery of the retina, the ciliary marginal zone (CMZ), produce new neurons in the retina throughout life.” (Perron and Harris, 2000); “The retina of many fish and amphibians grows throughout life, roughly matching the overall growth of the animal. The new retinal cells are continually added at the anterior margin of the retina, in a circumferential zone of cells” (Kubota et al., 2002); “The retinas of lower vertebrates grow throughout life from retinal stem cells (RSCs) and retinal progenitor cells (RPCs) at the rim of the retina.” (Wan et al., 2016); “In the retina of teleost fish, cell addition continues throughout life involving proliferation and axonal growth.” (García-Pradas et al., 2018) to name a few. However, studies from our group in the sea lamprey, Petromyzon marinus, and the catshark, *Scyliorhinus canicula*, revealed the loss of proliferative activity in the retina of adult individuals of these ancient vertebrate groups (Villar-Cheda et al., 2008; Hernández-Núñez et al., 2021). This raised the possibility that a continuous proliferative activity throughout life in the retina was a derived characteristic of modern teleost fishes and not the ancestral character common to all fish groups (Hernández-Núñez et al., 2021).

Based on our recent work in sharks (Hernández-Núñez et al., 2021), we decided to revisit the teleost literature on this topic (see Table S1). We observed that assumptions about a continuous proliferative and neurogenic activity in the retina of teleost fishes are mainly supported by work on juveniles and young adults, by work using animals in which the precise age is not indicated or by studies that did not systematically quantify the proliferative activity at different developmental and postnatal stages (see Table S1). In the zebrafish, *Danio rerio*, qualitative assessments of proliferating cells labelled with bromo-deoxyuridine (BrdU) in the peripheral and central retina from embryonic [24 hours post-fertilization (hpf)] to young adult (6-8 months) stages, revealed a sharp decline in the rate of retinal growth between the 3 and 4 days post-fertilization (dpf) and a decline in the rate of cell addition between embryos and young adults (Marcus et al., 1999). More recently, a thorough study by Van Houcke et al. (2019; see Table S1) evaluated the relative contribution of cellular addition and tissue stretching to retinal growth in the adult zebrafish from 6 to 48 months post-fertilization (mpf). By using immunohistochemical staining for proliferating cell nuclear antigen (PCNA) they quantified progenitor cell proliferation in the adult CMZ and found that the neurogenic capacity of the CMZ strongly declines between 6 and 12 months of age and continues at very low rates up to 48 months post-fertilization (mpf), though new cells continue to be added to the retina throughout life. However, while proliferating cells were also detected in the central retina, cell proliferation was not investigated over time in this area. On the other hand, since PCNA expression can also be detected long after cell cycle exit and can also indicate DNA repair or cell death (Mandyam et al., 2007 and at references therein), using PCNA alone as a proliferation marker may overestimate the number of cells progressing through the cell cycle, specially at the oldest stages.

To address these issues, we systematically quantified both the number of cells progressing through the cell cycle (immunoreactive to PCNA) and the number of cells undergoing mitosis (immunoreactive to the mitosis marker phosphohistone H3; pH3) in the peripheral and central retina of zebrafish, covering all major life stages from early developing (2 dpf) to sexual maturation (1.5 to 3 mpf) and ageing [up to 3-4 years post-fertilization (ypf)]. Our results show that there is a progressive loss of proliferative activity in the retina throughout life in zebrafish, even despite the occurrence of a wave of secondary proliferation that occurs during sexual maturation. Importantly, mitotic activity is virtually absent in the retina of old animals.

## 2. Results

The zebrafish retina exhibits the typical morphology and structure of the vertebrate retina, with a CMZ located at the retinal margin containing different types of progenitor cells and a highly organized central retina (Fig. 1A), which can be observed from 2.5 dpf (Malicki et al., 1996), formed by three nuclear layers: the outer nuclear layer (ONL), where the nuclei of photoreceptors are located; the INL, where the nuclei of horizontal, bipolar, amacrine and Müller glia cells are located; and the ganglion cell layer (GCL), which contains the nuclei of ganglion cells. These cells become connected within two plexiform layers: the outer plexiform layer (OPL) and the inner plexiform layer (IPL).

**Figure 1.**
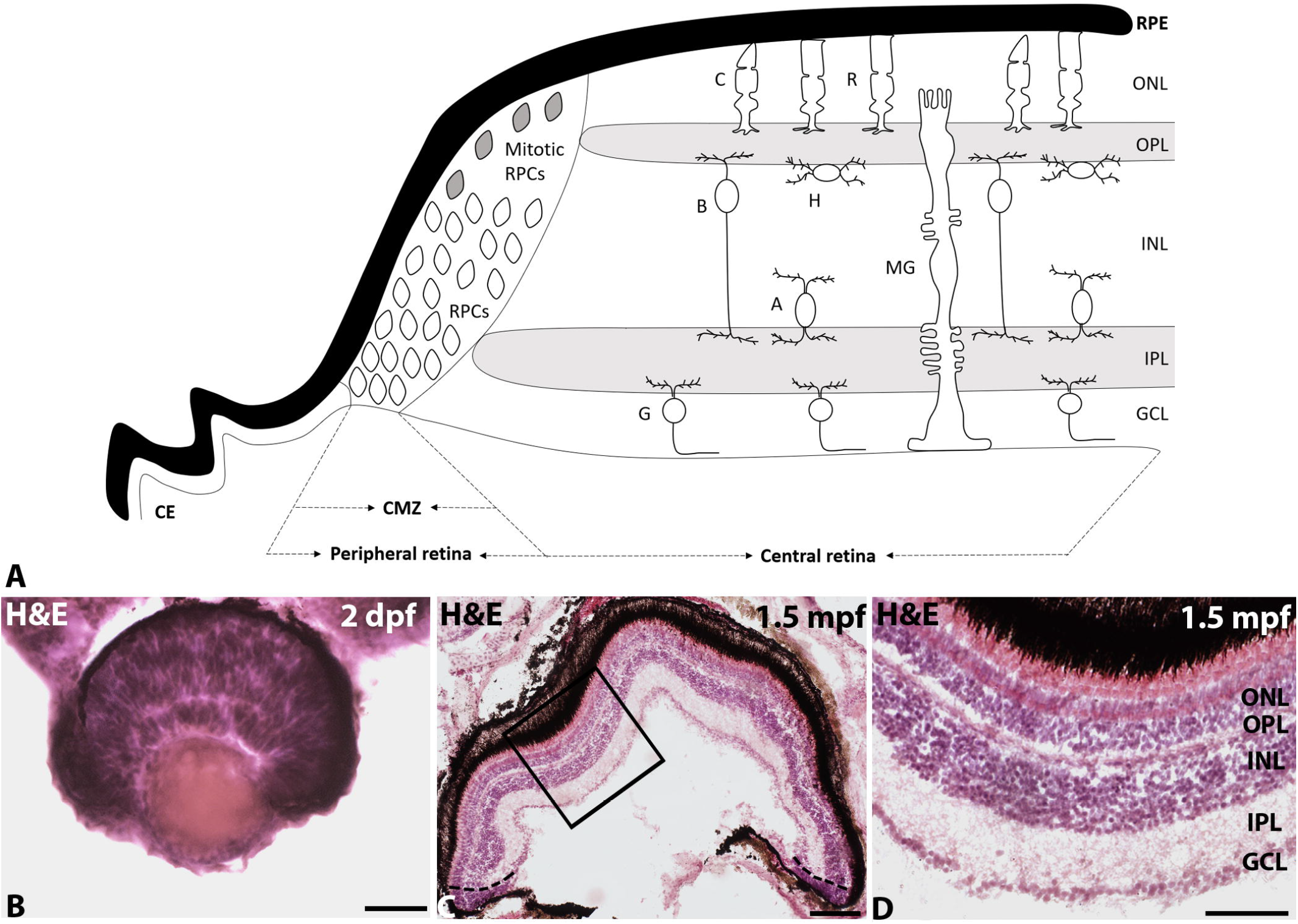
**A**. Schematic drawing of the mature zebrafish retina showing the retinal pigment epithelium (RPE) and the neural retina with two differentiated regions: the peripheral retina, which contains different types of progenitor cells (RPCs), and the central retina with a layered structure, which contains the outer (OPL) and inner (IPL) plexiform layers and three cell layers [ONL with the nuclei of cone (C) and rod (R) photoreceptors; inner nuclear layer (INL) with the nuclei of bipolar (B), amacrine (A), horizontal (H) and Müller glia (MG) cells; and a GCL with the nuclei of ganglion cells (G)]. B-D. Hemaetoxylin-eosin-stained transverse sections of the retina of 2 dpf (B) and 1.5 mpf (C, D) zebrafish showing the maturation of retinal organization. The dotted lines in C indicate the limit between the peripheral and the central retina. Figure D shows a detail of the central retina squared in figure C. Abbreviations: CE: ciliary epithelium; CMZ: ciliary marginal zone. Scale bars: B: 50 µm; C: 200 µm; D: 100 µm.

In this study, we analysed zebrafish of different developmental/life stages: 2 dpf, 4 dpf, 7 dpf, 1.5 mpf, 2.5 mpf, 3 mpf, 8.5 mpf, 18-20 mpf and 3-4 ypf. The 2 to 7 dpf period coincides with early zebrafish development (the zebrafish has a functional retina at about 3 dpf; Easter and Nicola, 1996). From 1.5 mpf to 3 mpf, zebrafish are in the process of sexual maturation, and at 8.5 mpf they are sexually mature and in peak fertility. From 18 mpf synaptic integrity begins to decline and a gradual increase of the senescence-associated β-galactosidase is observed in the RPE. This senescence marker is detected in the neural retina at 48 mpf (Van Houcke et al., 2019).

In 2 dpf animals, the retina did not show the typical layered organization of a mature vertebrate retina and it is mainly formed by neuroepithelial cells (Fig. 1B). As it can be observed in haematoxylin-eosin stained sections, the cell nucleus occupies almost the entire cell body in most retinal cells, which is characteristic of proliferating tissues (Fig. 1B). From 4 dpf, we observed the typical layered organization of the central retinal and that the CMZ was progressively reduced and restricted to the most marginal region with ageing (Figs. 1B-D).

### 2.1 Changes in proliferative activity with ageing

To compare the number of cells progressing through the cell cycle and cells undergoing mitosis in the zebrafish retina at different developmental/life stages we used double immunofluorescence to detect the expression of PCNA, which is present in proliferating cells during every phase of the cell cycle, peaking from G1 to S and decreasing at G2/M, and pH3, a marker of M-phase cells (Zerjatke et al., 2017). Our combinatorial analysis by studying both PCNA and pH3 expression helps to overcome the problem of using PCNA alone as a proliferation marker, since the latter can be also indicative of DNA repair or cell death (see introduction), which can lead to an overestimation of the number of proliferating cells. The number of PCNA+ or pH3+ cells is given as the mean number of cells per retinal section for each animal to allow for a comparison between specimens of different age and size.

Mitotic (pH3+) cells were mainly observed in the peripheral retina and in this region most of them were located in the apical surface, i.e. near the ventricle (Figs. 2A-C). However, some ectopic mitoses were also observed in the different layers of the central retina at the different developmental/life stages (Figs. 2A, C). pH3+ cells were almost absent in the whole retina of aged specimens (Fig. 2D). Most of the cells progressing through the cell cycle (PCNA+) were also located in the peripheral retina (Figs. 2A-D). However, as for pH3+ cells, PCNA+ cells were also present in the different cell layers of the central retina (Fig 2C). The number of PCNA+ cells was highly reduced in the whole retina of aged specimens (Fig. 2D).

**Figure 2.**
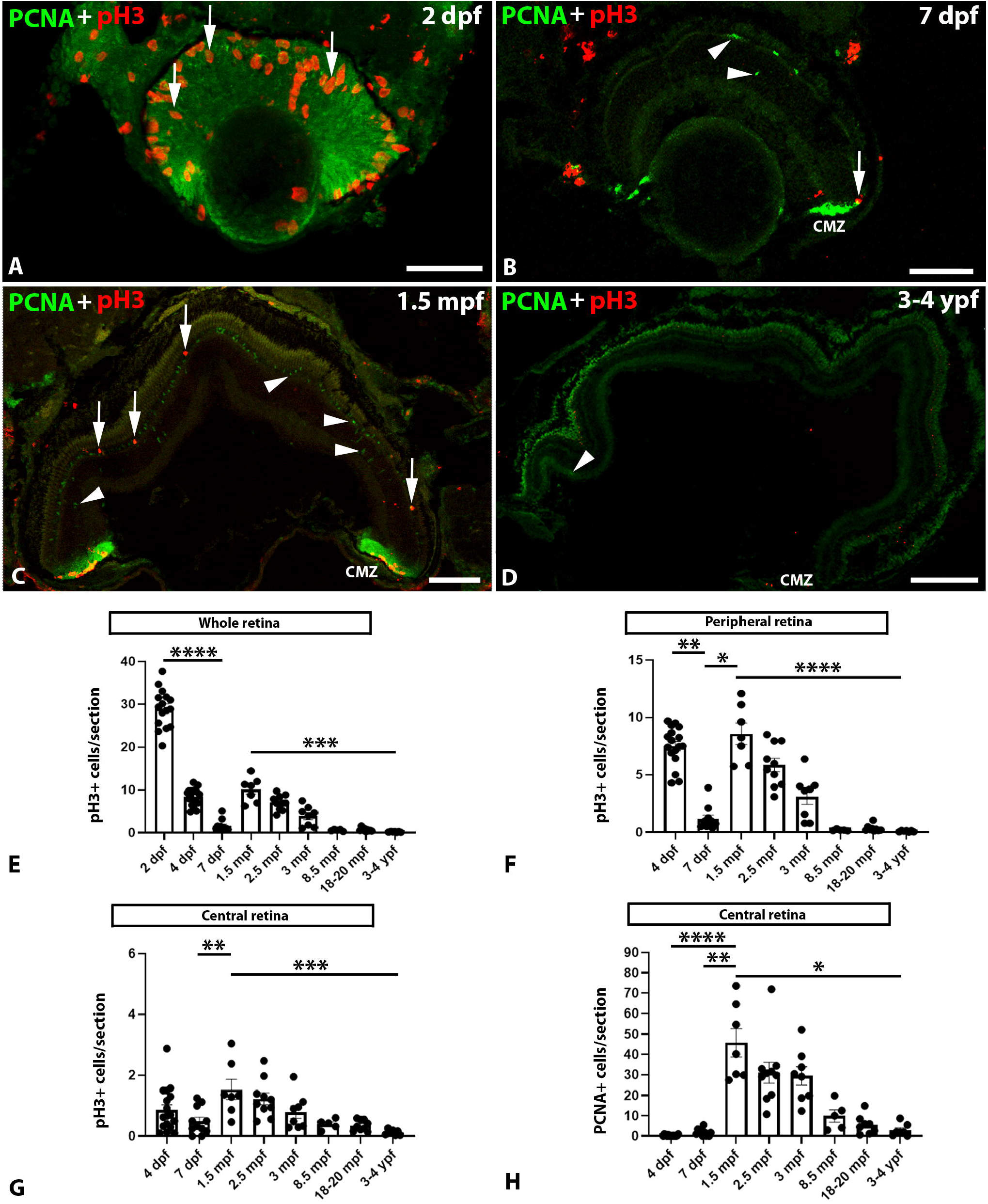
**A-D**. Transverse sections of the retina of 2 dpf (A), 7 dpf (B), 1.5 mpf (C) and 3-4 ypf (D) zebrafish specimens showing the presence of PCNA (arrowheads) or pH3 (arrows) positive cells. In 2 dpf to 1.5 mpf zebrafish (A-C) PCNA or pH3 expressing cells were mainly located in the peripheral retina (CMZ) but also in the central retina, while in 3-4 ypf specimens pH3+ and PCNA+ cells almost disappeared (D). Scale bars: A, B: 50 µm; C: 100 µm; D: 200 µm. E. Graph showing significant changes in the number of pH3+ cells/section in the whole retina at different ages (Kruskal-Wallis test, p < 0.0001). F. Graph showing significant changes in the number of pH3+ cells/section in the peripheral retina at different ages (Kruskal-Wallis test, p < 0.0001). G. Graph showing significant changes in the number of pH3+ cells/section in the central retina at different ages (one-way ANOVA, p < 0.0001). H. Graph showing significant changes in the number of PCNA+ cells/section in the central retina at different ages (Kruskal-Wallis test, p < 0.0001). Mean ± S.E.M. data and data on statistical multiple comparisons related to these graphs can be found on File S1. Asterisks indicate different levels of statistical significance: *, p < 0.05; **, p < 0.01; ***, p < 0.001; ****, p < 0.0001.

In the whole retina, the number of pH3+ cells significantly and progressively decreased during early development from 2 to 7 dpf (Fig. 2E; File S1). Interestingly, we observed an increase in mitotic activity in the whole retina at 1.5 mpf (which did not reach levels of 2 dpf animals; Fig. 2E; File S1). From 1.5 mpf onwards, we observed a significant and progressive decline in the number of mitotic cells with ageing (Fig. 2E; File S1). Importantly, mitotic cells were almost absent in the retina of aged specimens (from 8.5 mpf to 3-4 ypf, Fig. 2E; File S1).

Very similar trends in temporal expression patterns were observed when looking separately at the number of pH3+ cells per section in the CMZ (peripheral; Fig. 2F; File S1) and central (Fig. 2G; File S1) retinas or at the number of PCNA+ cells per section in the central retina (Fig. 2H; File S1).

Our results indicate that proliferative activity decreases with age in both the peripheral and central regions of the zebrafish retina, despite the occurrence of a secondary wave of proliferation during sexual maturation (i.e. 1.5 to 3 mpf). Importantly, mitotic activity is virtually absent in aged specimens.

### 2.2 Changes in the location of proliferating cells of the central retina with age

We quantified separately the number of pH3+ and PCNA+ cells in the cell layers of the central retina (GCL, INL and ONL) at the different developmental and life stages (Fig. 3; File S2). Interestingly, in early developing 4 dpf specimens the number of pH3+ and PCNA+ cells was significantly higher in the INL than in the GCL or ONL (Fig. 3; File S2), whereas from 7 dpf to 18-20 mpf (both included) the number of pH3+ and PCNA+ cells was significantly higher in the ONL than in GCL or INL (Fig. 3; File S2). As can be observed in Figure 3, this difference is highly significant during sexual maturation (1.5 to 3 mpf). In the oldest animals (3 to 4 ypf) the very few pH3+ or PCNA+ cells located in the central retina did not show differential distribution in the cell layers (Fig. 3; File S2). These results suggest that the progenitor cells that remain in the mature central retina after early development could mainly contribute to the production of mature cell types of the ONL.

**Figure 3.**
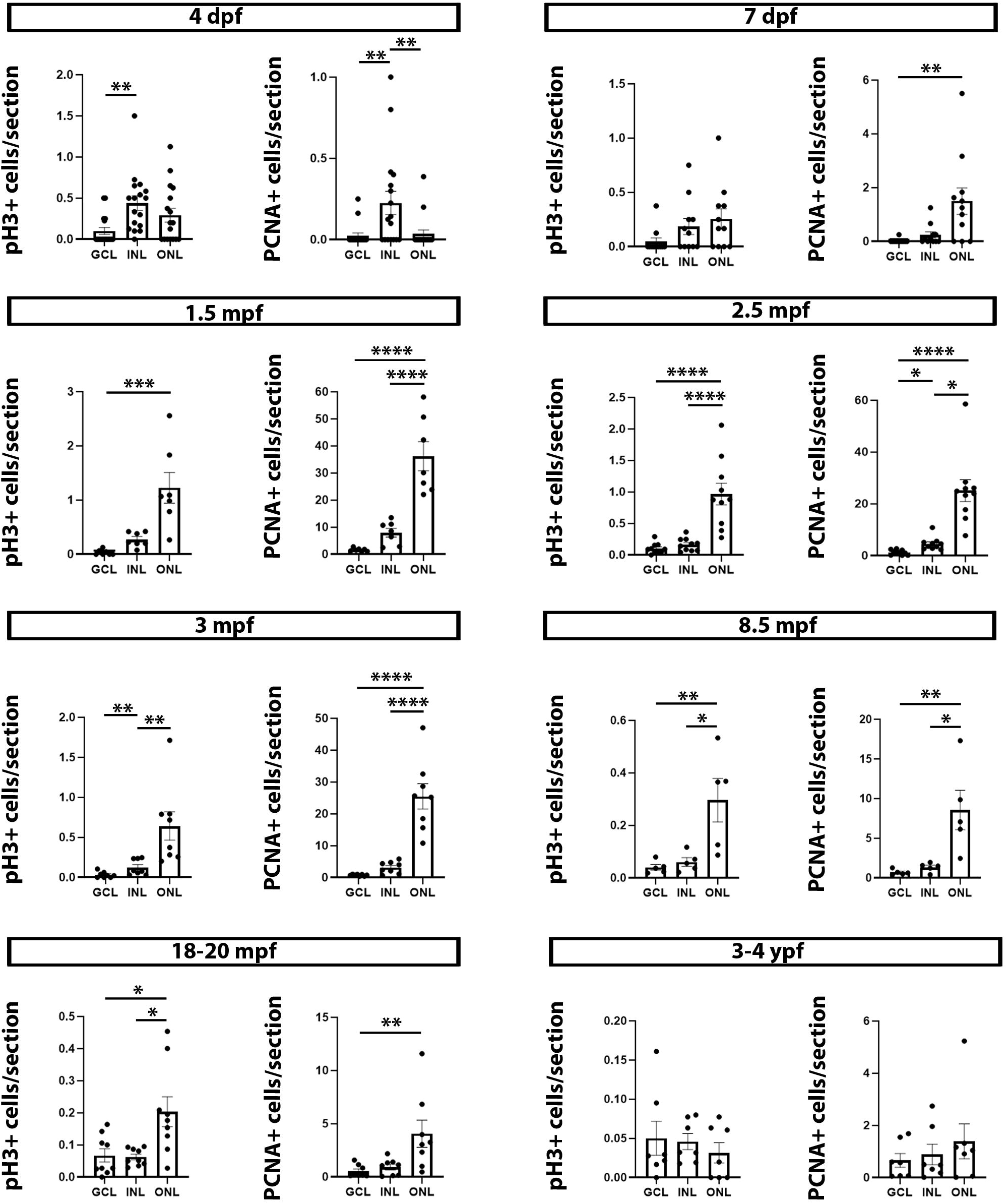
Graphs showing the differential distribution of pH3+ and PCNA+ cells in cell layers of the central retina at different developmental and life stages. 4 dpf specimens: pH3 (Kruskal-Wallis test, p = 0.0022), PCNA (Kruskal-Wallis test, p = 0.0018). 7 dpf specimens: pH3 (Kruskal-Wallis test, p = 0.0655), PCNA (Kruskal-Wallis test, p = 0.0024). 1.5 mpf specimens: pH3 (Kruskal-Wallis test, p < 0.0001), PCNA (one-way ANOVA, p < 0.0001). 2.5 mpf specimens: pH3 (one-way ANOVA, p < 0.0001), PCNA (Kruskal-Wallis test, p < 0.0001). 3 mpf specimens: pH3 (one-way ANOVA, p = 0.0009), PCNA (one-way ANOVA, p < 0.0001). 8.5 mpf specimens: pH3 (one-way ANOVA, p = 0.0055), PCNA (one-way ANOVA, p = 0.0038). 18-20 mpf specimens: pH3 (Kruskal-Wallis test, p = 0.0121), PCNA (Kruskal-Wallis test, p = 0.0084). 3-4 ypf specimens: pH3 (Kruskal-Wallis test, p = 0.4814), PCNA (Kruskal-Wallis test, p = 0.8584). Mean ± S.E.M. data and data on statistical multiple comparisons related to these graphs can be found on File S2. Asterisks indicate different levels of statistical significance: *, p < 0.05; **, p < 0.01; ***, p < 0.001; ****, p < 0.0001.

## 3. Discussion

As indicated in the introduction, it is largely assumed that the retina of fishes shows continuous and active proliferation and neurogenesis throughout life. This assumption is based on previous work in teleost models in which the presence of proliferating cells was only studied in juveniles or young adults, in animals in which the precise age was not defined or known by the authors of the study or without perform-ing quantitative comparisons between all life stages or ages (see Table S1). This feature of throughout life neurogenesis does not apply to lampreys or cartilaginous fishes, in which proliferative activity is virtually absent in adult animals (Villar-Cheda et al., 2008; Hernández-Núñez et al., 2021). Moreover, some of the previous studies on teleost fishes provided qualitative descriptions that also suggested a loss of proliferating cells with age (see Table S1). For example, Johns and Fernald (1981) reported that, when studying African cichlid and goldfish juveniles and adults, dividing cells in the ONL were easier to demonstrate in younger fish. In zebrafish, Marcus et al. (1999) also indicated that the number of BrdU labelled cells was greater in the peripheral and central retina of embryos than in young adults (6-8 mpf). A recent study by Van Houcke et al. (2019) showed a decline in cell proliferation (PCNA+ cells) in the zebrafish CMZ from 6 to 48 mpf. However, proliferation within the central retina of zebrafish was not quantified over time. Besides, assessment of progenitor cell proliferation relied only on PCNA expression, despite its use can lead to overestimation of proliferation in aged animals (see introduction).

Here, we obtained quantitative data comparing cell cycle progression (PCNA+ cells/section) and mitotic activity (pH3+ cells/section) in both the CMZ and the central zebrafish retina at different ages and covering all major life stages. Our results show that there is a drastic decline in proliferative activity from 2 dpf to 7 dpf, a continuous reduction in the number of proliferating cells in sexually maturing and old animals, and that cells undergoing mitosis are virtually absent in old animals. This is in good agreement with previous reports of a drastic decrease in cell proliferation in early larvae (between the 3 dpf and 4 dpf; Marcus et al., 1999) and with reports of a significant proliferation decrease in early adulthood (6-12 mpf), with very reduced proliferation rates a at mid (18-24 mpf) and late (36-38 mpf) adult stages (Van Houcke et al., 2019). As expected, the number of PCNA+ cells reported in the CMZ by Van Houcke et al. (2019) was higher to that of pH3+ cells in this region at similar life stages (present results), since the latter are only a fraction of the number of cells progressing through the cell cycle. Our quantitative results in zebrafish reveal a similar pattern to that reported in our work in lampreys and sharks showing a loss of proliferating and mitotic cells in the adult retina (Villar-Cheda et al., 2008; Hernández-Núñez et al., 2021). However, it seems that retinal proliferative activity is maintained at a higher rate in adult zebrafish than in adult lampreys (no PCNA+ cells; Villar-Cheda et al., 2008) or sharks (very few PCNA+ cells and almost absent pH3+ cells; Hernández-Núñez et al., 2021).

Unlike previous reports on proliferation in the zebrafish retina, our systematic analysis allowed unveiling the occurrence of a secondary wave of proliferation during sexual maturation (i.e. from 1.5 to 3 mpf) affecting both the CMZ and the central retina. Only a study by Bernardos et al. (2007) provided a qualitative description indicating that BrdU+ cells in the ONL were observed in higher numbers in 1 to 2 mpf animals than in 7 dpf animals. This increase in cell proliferation at 1.5 mpf (present results) did not reach levels of the early developing (2 dpf) period but it was significantly higher than in 7 dpf specimens. This secondary wave of proliferation could be related to an earlier peak of cell death that occurs in the retina (especially in the ONL) of 7 dpf zebrafish (Biehlmaier et al., 2001). This increase in cell proliferation could allow to replace cells lost during this critical period in which fish transition from acquiring nutrition from their yolk to active feeding. This secondary wave of proliferation could be also related to retinal adaptations that might be needed for sexual behaviors, especially since the integration of multisensory information between olfaction and vision has been implicated in mating-like behaviors in zebrafish (Li et al., 2017). However, current data has only implicated dopaminergic interplexiform and retinal ganglion cells in this olfactovisual centrifugal pathway (Li et al., 2017), which would not explain why proliferation and neurogenesis are needed in the ONL (see below). Future studies should decipher whether this secondary wave of proliferation is only needed to replace lost retinal cells or if it is related to retinal adaptations needed for sexual (or adult) behaviors in zebrafish.

By looking at the distribution of proliferating/mitotic cells in the cell layers of the central retina at different ages we observed that in early developing 4 dpf specimens the number of pH3+ and PCNA+ cells was higher in the INL, whereas in older animals they were more abundant in the ONL. Previous studies have shown that progenitor cells of the central retina (Müller glia) in juvenile/adult goldfish (Johns and Fernald, 1981; Johns, 1982; Otteson et al., 2001) and in juvenile zebrafish (Bernardos et al., 2007; Morris et al., 2008a see Morris et al., 2008b; Lenkowski and Raymond, 2014) generate new rods, which indicates that the higher cell proliferation and mitotic activity we observed in the ONL of juvenile and adult zebrafish is related to rod generation. As far as we are aware the generation of other retinal cell types from INL and ONL progenitors of the uninjured juvenile/adult teleost retina has not been reported, though injury-induced proliferating Müller glial cells can regenerate all retinal cell types, including cones (Bernardos et al., 2007; Morris et al, 2008a, b; Lenkowski and Raymond, 2014), which has led to suggest that the neuronal progenitors produced by Müller glia are multipotent and can revert to an earlier lineage under the influence of certain microenvironmental signals (Bernardos et al., 2007, Morris et al., 2008; reviewed in Morris et al., 2008b; Stenkamp, 2011). The zebrafish retina presents 5 main types of photoreceptors (4 cones and 1 rod) and the 5 types of photoreceptors are generated during early development (Crespo and Knust, 2018). Perhaps one or more of these photoreceptor types are specifically needed for mating/courtship/adult behaviors and could be generated in extra numbers during sexual maturation, which could explain the secondary wave of cell proliferation we detected in zebrafish juveniles. Since during courtship and spawning female zebrafish discriminate between sexes using visual cues in which the male yellow coloration is critical (Hutter et al., 2012), it is tempting to hypothesize that specific cones might be needed at this life stage. However, to our knowledge, microenviromental signals other than retinal injury driving cone generation from progenitors in the central retina have not been experimentally assessed. Future work should attempt to study whether cones could be also generated from these dividing progenitor cells of the central retina, especially during the previously undetected secondary wave of proliferation at the time of sexual maturation.

## 4. Material and Methods

### 4.1. Animals

Two days post-fertilization (2 dpf, n = 16), 4 dpf (n = 17), 7 dpf (n = 11), 1.5 mpf (n = 7), 2.5 mpf (n = 10), 3 mpf (n = 8), 8.5 mpf (n = 5), 18-20 mpf (n = 9) and 3-4 years post-fertilization (ypf, n = 7) zebrafish (*Danio rerio*) specimens were used in this study. Zebrafish were kept in aquaria under standard conditions of temperature (28 °C), light cycle (14 h of light and 10 h of darkness) and pH (7.0) until use for experimental procedures. All juvenile/adult fish were kept at a density of 3 fish/litre and food supply was under strict standardized control to avoid differences in animal size in each of the experimental groups (Van Houcke et al., 2019).

### 4.2. Tissue preparation for histology

Animals were deeply anesthetized with 0.0016% tricaine methanesulfonate (MS-222, Sigma-Aldrich, St. Louis, MO), euthanized and fixed by immersion in 4% paraformaldehyde in 0.1M phosphate-buffered saline pH 7.4 (PBS) for 2 h (from 2 dpf to 7 dpf) and 1 day (from 1.5 mpf to 3-4 ypf) at 4 °C. After fixation, the lens was removed from the eye in specimens from 1.5 mpf onwards. Eyes were dissected out from the rest of the body in specimens from 8.5 mpf onwards. After rinsing in PBS, the animals or eyes were cryoprotected with 30 % sucrose in PBS, embedded in Neg-50TM (Thermo Scientific, Kalamazoo, MI) and frozen with liquid nitrogen-cooled isopentane. Transverse sections (18 μm thick) were obtained on a cryostat and mounted on Superfrost Plus slides (Menzel-Glasser, Madison, WI).

### 4.3. Haematoxylin-eosin staining

Some sections from 2 dpf and 1.5 mpf specimens were stained with haematoxylin-eosin following standard protocols. Briefly, cryostat sections were dried at room temperature (RT), rinsed in 0.05 M Tris-buffered (pH 7.4) saline (TBS) for 10 min and stained with haematoxylin solution for 10 min. Sections were subsequently rinsed in tap water until removal of the excess of haematoxylin, in distilled water for 10 min and then stained with eosin for 2 min. Finally, the sections were dehydrated and mounted in DPX mounting medium (Scharlau, Sentmenat, Spain).

### 4.4. Immunofluorescence

Sections were first pre-treated with 0.01 M citrate buffer pH 6.0 for 30 min at 90 °C for heat-induced epitope retrieval, allowed to cool for 20 min at RT and rinsed in TBS for 5 min. Then, sections were incubated overnight at RT with a combination of two primary antibodies: a mouse monoclonal anti-PCNA (1:500; Sigma-Aldrich; catalogue number P8825; RRID: AB_477413) and a rabbit polyclonal anti-pH3) (1:300; Millipore; Billerica, MA; catalogue number 06-570; RRID: AB_310177). Sections were rinsed 3 times in TBS for 10 min each and incubated for 1 h at RT with a combination of fluorescent dye-labelled secondary antibodies: a Cy3-conjugated goat anti-rabbit (1:200; Invitrogen, Waltham, MA. USA; catalogue number A10520) and a FITC-conjugated goat anti-mouse (1:200; Invitrogen; catalogue number F2761). All antibody dilutions were made in TBS containing 15 % normal goat serum (Millipore), 0.2 % Triton X-100 (Sigma-Aldrich) and 2 % BSA (Sigma-Aldrich). Sections were then rinsed 3 times in TBS for 10 min each and in distilled water for 30 min, allowed to dry for 30 min at 37 °C and mounted in MOWIOL® 4-88 (Calbiochem, Darmstadt, Ger-many).

### 4.5. Specificity of antibodies

Anti-PCNA antibodies (including the one used in this study) have been traditionally used to label proliferating cells in the retina of different fish (including zebrafish): *Oryzias latipes* (Negishi et al., 1990); *Haplochromis burtoni* (Mack and Fernald, 1995, 1997); *Onchorynchus mykiss* (Julian et al., 1998); *Tinca tinca* (Velasco et al., 2001; Cid et al., 2002; Jimeno et al., 2003); *Carasius auratus* (Negishi et al., 1990; Cid et al., 2002); *Salmo trutta fario* (Candal et al., 2005); *Danio rerio* (Cid et al., 2002; Amini et al., 2019); *Scyliorhinus canicula* (Ferreiro-Galve et al., 2010a, b; 2012; Bejarano-Escobar et al., 2012; Sánchez-Farías and Candal, 2015; 2016; Hernández-Núñez et al., 2021); and *Petromyzon marinus* (Villar-Cheda et al., 2008). Anti-pH3 antibodies (including the one used in this study) have been also commonly used to label mitotic cells in the retina of fish (including zebrafish): *Carasius auratus* (Otteson et al., 2001); *Scyliorhinus canicula* (Ferreiro-Galve et al., 2010a; Bejarano-Escobar et al., 2012; Hernández-Núñez et al., 2021); and *Danio rerio* (Jensen et al., 2001; Godinho et al., 2007; Weber et al., 2014).

### 4.6. Image acquisition

Brightfield images of haematoxylin-eosin stained sections were taken with an Olympus BX51 microscope equipped with an Olympus DP71 camera. Images of fluorescent labelled sections were taken with a Leica TCS-SP2 confocal microscope with a combination of blue and green excitation lasers. Confocal optical sections were taken at steps of 1 μm along the z-axis. Collapsed images of the whole retinal sections (18 µm) were obtained with the LITE software (Leica, Wetzlar, Germany). For figure preparation, contrast and brightness of the images were minimally adjusted using Adobe Photoshop CS4 (Adobe, San Jose, CA).

### 4.7. Cell quantifications and statistical analyses

We quantified the number of mitotic cells (pH3+) in the whole retina (peripheral and central retina) and cells progressing through the cell cycle (PCNA+) in the central retina. The number of PCNA+ cells was not quantified in the peripheral retina because at some stages (mainly 2 dpf) the high number of positive cells impeded to clearly differentiate between cells individually. In any case, the number of PCNA+ cells/section in the CMZ of adult zebrafish (6 to 48 mpf) was previously examined by Van Houcke et al. (2019).

The number of pH3+ and PCNA+ cells was manually counted under a fluorescence microscope in one out of each two consecutive retinal sections (thickness of 18 μm). The limit between the peripheral (CMZ) and the central retina was established based on the expression pattern of PCNA, which is mainly found in the most peripheral region of the retina. In 2 dpf specimens, we did not separate pH3+ cell quantifications of the peripheral and central retinas because at this developmental stage PCNA was highly expressed throughout the entire retina, which impeded to establish a clear limit between both regions. We calculated the mean number of cells per section for each retina. Then, we calculated the mean number of cells per retinal section for each animal based on the values for the two retinas and used that value for statistical analyses (each dot in the graph represents one animal). We also quantified the differential distribution of pH3+ and PCNA+ cells in the different cell layers of the central retina from 4 dpf onwards: the GCL, the INL and the ONL.

Statistical analyses were performed with Prism 8 (GraphPad software, La Jolla, CA). Normality of the data was determined with the Kolmogorov-Smirnov test. An ordinary one-way ANOVA followed by a Turkey’s multiple comparison test was used to determine statistically significant differences in normally distributed data. A Kruskal-Wallis test followed by a Dunn’s multiple comparison test was used to determine statistically significant differences in non-normally distributed data.

## 5. Conclusions

Our results reveal a decline in constitutive proliferative activity in the zebrafish retina with ageing. Importantly, mitotic activity is virtually absent in aged animals. Statements regarding continuous and high proliferative activity in the fish (teleost) retina throughout life should be nuanced.

Interestingly, our systematic study also detected the presence of a secondary wave of cell proliferation during sexual maturation in the zebrafish retina. A possible relationship to the generation of photoreceptors needed for sexual/adult behaviours is suggested. However, future work should try to decipher the origin and destiny of these progenitor cells and their relationship to zebrafish behaviour days post-fertilization (2 dpf, n = 16), 4 dpf (n = 17), 7 dpf (n = 11), 1.5 months post-fertilization (mpf, n = 5), 2.5 mpf (n = 10), 3 mpf (n = 8), 8.5 mpf (n = 5), 18-20 mpf (n = 9) and 3-4 years post-fertilization (ypf, n= 5) zebrafish (*Danio rerio*) specimens were used in this study. Zebrafish were kept in aquaria under standard conditions of temperature (28 °C), light cycle (14 h of light and 10 h of darkness) and pH (7.0) until use for experimental procedures. All experimental procedures were performed according to the regulations and laws established by the European Union (2010/63/UE) and by the Spanish Royal Decree 1386/2018 for the care and handling of animals in research and were approved by the Bioethics Committee of the University of Santiago de Compostela.

## Supplementary Materials

**Table S1**. Studies demonstrating the presence of proliferating cells in the juvenile/adult retina of different teleost species.

Only studies that included juvenile/adult specimens in their analyses were added to the table. Studies looking at cell proliferation during regenerative processes were not included. Note that most of the studies only looked at a specific stage/age/size and that these were usually juveniles or young adults. Sometimes the authors only referred to the animals as “adults” without any indication or age or size. In most of the few cases in which several life stages were compared no systematic quantifications or statistical analyses were performed. Only the recent study by Van Houcke et al. (2019) quantified cell proliferation (PCNA) in the CMZ of adult zebrafish of different ages (6 to 48 mpf). Moreover, only one study in goldfish juveniles used a marker of mitotic cells (pH3; Otteson et al., 2001).

**File S1**. Mean ± S.E.M. data and data on statistical multiple comparisons related to graphs shown in Figure 2.

**File S2**. Mean ± S.E.M. data and data on statistical multiple comparisons related to graphs shown in Figure 3.

## Author Contributions

Conceptualization, IH-N, FA, EC and AB-I; methodology, IH-N, AQ-R; formal analysis, IH-N, FA, EC, AB-I; resources, LS, EC and AB-I; writing—original draft preparation, IH-N, FA, EC, AB-I; writing—review and editing, IH-N, AQ-R, LS, FA, EC, AB-I; funding acquisition, IH-N, EC, AB-I. All authors have read and agreed to the published version of the manuscript.

## Institutional Review Board Statement

The study was conducted according to the regulations and laws established by the European Union (2010/63/UE) and by the Spanish Royal Decree 1386/2018 for the care and handling of animals in research and were approved by the Bioethics Committee of the University of Santiago de Compostela and the Xunta de Galicia (reference MR 110250).

## Data Availability Statement

Data are contained within the article and materials can be requested from the authors upon reasonable request.

## References

1. Alunni, A.; Bally-Cuif, L. A comparative view of regenerative neurogenesis in vertebrates. Development. 2016, 143(5), 741–753. doi: 10.1242/dev.122796.

2. Álvarez-Buylla, A.; Lim, D.A. For the long run: maintaining germinal niches in the adult brain. Neuron. 2004, 41, 683–686. doi: 10.1016/S0896-6273(04)00111-4.

3. Amato, M.A.; Arnault, E.; Perron, M. Retinal stem cells in vertebrates: Parallels and divergences. Int. J. Dev. Biol. 2004, 48, 993–1001. doi: 10.1387/ijdb.041879ma.

4. Amini, R.; Labudina, A.A.; Norden, C. Stochastic single cell migration leads to robust horizontal cell layer formation in the vertebrate retina. Development. 2019, 146(12):dev173450. doi: 10.1242/dev.173450.

5. Bejarano-Escobar, R.; Blasco, M.; Durán, A.C.; Rodríguez, C.; Martín-Partido, G.; Francisco-Morcillo, J. Retinal histogenesis and cell differentiation in an elasmobranch species, the small-spotted catshark Scyliorhinus canicula. J. Anat. 2012, 220, 318–335. doi: 10.1111/j.1469-7580.2012.01480.x.

6. Bélanger, M.C.; Robert, B.; Cayouette, M. Msx1-positive progenitors in the retinal ciliary margin give rise to both neural and non-neural progenies in mammals. Dev. Cell. 2017, 40(2), 137–150. doi: 10.1016/j.devcel.2016.11.020.

7. Bernardos, R.L.; Barthel, L.K.; Meyers, J.R.; Raymond, P.A. Late-stage neuronal progenitors in the retina are radial Müller glia that function as retinal stem cells. J. Neurosci. 2007, 27(26), 7028–7040. doi: 10.1523/JNEUROSCI.1624-07.200.

8. Biehlmaier, O.; Neuhauss, S.C.; Kohler, K. Onset and time course of apoptosis in the developing zebrafish retina. Cell Tissue Res. 2001, 306(2), 199–207. doi: 10.1007/s004410100447.

9. Candal, E.; Anadón, R.; DeGrip, W.J.; Rodríguez-Moldes, I. Patterns of cell proliferation and cell death in the developing retina and optic tectum of the brown trout. Brain Res. Dev. 2005, 154(1), 101–119. doi: 10.1016/j.devbrainres.2004.10.008

10. Cid, E.; Velasco, A.; Ciudad, J.; Orfao, A.; Aijon, J.; Lara, J.M. Quantitative evaluation of the distribution of proliferating cells in the adult retina in three cyprinid species. Cell Tissue Res. 2002, 308, 47–59. doi: 10.1007/s00441-002-0529-8.

11. Crespo, C.; Knust, E. Characterisation of maturation of photoreceptor cell subtypes during zebrafish retinal development. Biol. Open. 2018, 7(11):bio036632. doi: 10.1242/bio.036632.

12. Das, A.V.; James, J.; Rahnenführer, J.; Thoreson, W.B.; Bhattacharya, S.; Zhao, X.; Ahmad, I. Retinal properties and potential of the adult mammalian ciliary epithelium stem cells. Vision Res. 2005, 45(13), 1653–1666. doi: 10.1016/j.visres.2004.12.017.

13. Das, A.V.; Zhao, X.; James, J.; Kim, M.; Cowan, K.H.; Ahmad, I. Neural stem cells in the adult ciliary epithelium express GFAP and are regulated by Wnt signaling. Biochem. Biophys. Res. Commun. 2006, 339(2), 708–716. doi: 10.1016/j.bbrc.2005.11.064.

14. Doetsch, F. A niche for adult neural stem cells. Curr. Opin. Genet. Dev. 2003, 13(5), 543–550. doi: 10.1016/j.gde.2003.08.012.

15. Engelhardt, M.; Bogdahn, U.; Aigner, L. Adult retinal pigment epithelium cells express neural progenitor properties and the neuronal precursor protein doublecortin. Brain Res. 2005, 1040(1-2), 98–111. doi: 10.1016/j.brainres.2005.01.075.

16. Easter, S.S., Jr.; Nicola, G.N. The development of vision in the zebrafish (Danio rerio). Dev. Biol. 1996, 180, 646–663. https://doi.org/10.1006/dbio.1996.0335

17. Eymann, J.; Salomies, L.; Macrì, S.; Di-Poï, N. Variations in the proliferative activity of the peripheral retina correlate with postnatal ocular growth in squamate reptiles. J. Comp. Neurol. 2019, 527, 2356–2370. doi: 10.1002/cne.24677.

18. Fausett, B.V.; Goldman, D. A role for α1 tubulin-expressing Müller glia in regeneration of the injured zebrafish retina. J. Neurosci. 2006, 26(23), 6303–6313. doi: 10.1523/JNEUROSCI.0332-06.2006.

19. Fernald, RD. Teleost vision: seeing while growing. J. Exp. Zool. 1991, Suppl. 5, 167–180. doi: 10.1002/jez.1402560521.

20. Ferreiro-Galve, S.; Rodríguez-Moldes, I.; Anadón, R.; Candal, E. Patterns of cell proliferation and rod photoreceptor differentiation in shark retinas. J. Chem. Neuroanat. 2010a, 39, 1–14. doi: 10.1016/j.jchemneu.2009.10.001.

21. Ferreiro-Galve, S.; Rodríguez-Moldes, I.; Candal, E. Calretinin immunoreactivity in the developing retina of sharks: comparison with cell proliferation and GABAergic system markers. Exp. Eye Res. 2010b, 91, 378–386. doi: 10.1016/j.exer.2010.06.011.

22. Ferreiro-Galve, S.; Rodríguez-Moldes, I.; Candal, E. Pax6 expression during retinogenesis in sharks: comparison with markers of cell proliferation and neuronal differentiation. J. Exp. Zool. B Mol. Dev. Evol. 2012, 318, 91–108. doi: 10.1002/jezb.21448.

23. Ferretti, P. Is there a relationship between adult neurogenesis and neuron generation following injury across evolution? Eur. J. Neurosci. 2011, 34(6), 951–962. doi: 10.1111/j.1460-9568.2011.07833.x.

24. Fischer, A.J.; Reh, T.A. Transdifferentiation of pigmented epithelial cells: a source of retinal stem cells? Dev. Neurosci. 2001, 23(4-5), 268–276. doi: 10.1159/000048710.

25. Fischer, A.J; Reh, T.A. Growth factors induce neurogenesis in the ciliary body. Dev. Biol. 2003, 259(2), 225–240. doi: 10.1016/s0012-1606(03)00178-7.

26. Fischer, A.J.; Bosse, J.L.; El-Hodiri, M. The ciliary marginal zone (CMZ) in development and regeneration of the vertebrate eye. Exp. Eye Res. 2013, 116, 199–204. doi: 10.1016/j.exer.2013.08.018.

27. García-Pradas, L.; Gleiser, C.; Wizenmann, A.; Wolburg, H.; Mack, A.F. Glial cells in the fish retinal nerve fiber layer form tight junctions, separating and surrounding axons. Front. Mol. Neurosci. 2018, 11:367. doi: 10.3389/fnmol.2018.00367.

28. Godinho, L.; Williams, P.R.; Claassen, Y.; Provost, E.; Leach, S.D.; Kamermans, M.; Wong, R.O. Nonapical symmetric divisions underlie horizontal cell layer formation in the developing retina in vivo. Neuron. 2007, 56(4), 597–603. doi: 10.1016/j.neuron.2007.09.036.

29. Grandel, H.; Brand, M. Comparative aspects of adult neural stem cell activity in vertebrates. Dev Genes Evol. 2013, 223(1-2), 131–147. doi: 10.1007/s00427-012-0425-5.

30. Harris, W.A.; Perron, M. Molecular recapitulation: the growth of the vertebrate retina. Int. J. Dev. Biol. 1998, 42(3), 299–304.

31. He, J.; Zhang, G.; Almeida, A.D.; Cayouette, M.; Simons, B.D.; Harris, W.A. How variable clones build an invariant retina. Neuron. 2012, 75(5), 786–798. doi: 10.1016/j.neuron.2012.06.033.

32. Hernández-Núñez, I.; Robledo, D.; Mayeur, H.; Mazan, S.; Sánchez, L.; Adrio, F.; Barreiro-Iglesias, A.; Candal E. Loss of active neurogenesis in the adult shark retina. Front. Cell Dev. Biol. 2021, 9:628721. doi: 10.3389/fcell.2021.628721.

33. Hutter, S.; Hettyey, A.; Penn, D.J.; Zala, S.M. Ephemeral sexual dichromatism in zebrafish (Danio rerio). Ethology. 2012, 118, 1208–1218. doi: 10.1111/eth.12027.

34. Jensen, A.M.; Walker, C.; Westerfield, M. Mosaic eyes: a zebrafish gene required in pigmented epithelium for apical localization of retinal cell division and lamination. Development. 2001, 128(1), 95–105.

35. Jimeno, D.; Lillo, C.; Cid, E.; Aijón, J.; Velasco, A.; Lara, J.M. The degenerative and regenerative processes after the elimination of the proliferative peripheral retina of fish. Exp. Neurol. 2003, 179(2), 210–228. doi: 10.1016/S0014-4886(02)00020-1.

36. Johns, P.R. Formation of photoreceptors in larval and adult goldfish. J. Neurosci. 1982, 2(2), 178–198. doi: 10.1523/JNEUROSCI.02-02-00178.1982.

37. Johns, P.R.; Fernald, R.D. Genesis of rods in teleost fish retina. Nature. 1981, 293(5828), 141–142. doi: 10.1038/293141a0.

38. Julian, D.; Ennis, K.; Korenbrot, J.I. Birth and fate of proliferative cells in the inner nuclear layer of the mature fish retina. J. Comp. Neurol. 1998, 394(3), 271–282.

39. Kubota, R.; Hokoc, J.N.; Moshiri, A.; McGuire, C.; Reh, T.A. A comparative study of neurogenesis in the retinal ciliary marginal zone of homeothermic vertebrates. Dev. Brain Res. 2002, 134, 31–41. doi: 10.1016/S0165-3806(01)00287-5.

40. Kwan, J.W.; Lee M.J.; Mack A.F.; Chiu J.F.; Fernald R.D. Nonuniform distribution of cell proliferation in the adult teleost retina. Brain Res. 1996, 712, 40–44. doi: 10.1016/0006-8993(95)01426-8.

41. Lenkowski, J.R.; Raymond, P.A. Mu□ller glia: stem cells for generation and regeneration of retinal neurons in teleost fish. Prog. Retin. Eye Res. 2014, 40, 94–123. doi: 10.1016/j.preteyeres.2013.12.007.

42. Li, L.; Wojtowicz, J.L.; Malin, J.H.; Huang, T.; Lee, E.B.; Chen, Z. GnRH-mediated olfactory and visual inputs promote mating-like behaviors in male zebrafish. PLoS One. 2017, 12(3):e0174143. doi: 10.1371/journal.pone.0174143.

43. Lyall, A.H. The growth of the trout retina. Q. J. Micros. Sci. 1957, 98, 101–110.

44. Ma, R.T.Y.; Li, X.; Wang, S. Z. Reprogramming RPE to differentiate towards retinal neurons with Sox2. Stem Cells. 2009, 27, 1376–1387. doi: 10.1002/stem.48. doi: 10.1002/stem.48.

45. Mack, A.F.; Fernald, R.D. New rods move before differentiating in adult teleost retina. Dev. Biol. 1995, 170(1), 136–141. doi: 10.1006/dbio.1995.1202.

46. Mack, A.F.; Fernald, R.D. Cell movement and cell cycle dynamics in the retina of the adult teleost Haplochromis burtoni. J. Comp. Neurol. 1997, 388(3), 435–443.

47. Malicki, J.; Neuhauss, S.C.; Schier, A.F.; Solnica-Krezel, L.; Stemple, D.L.; Stainier, D.Y.; Abdelilah, S.; Zwartkruis, F.; Rangini, Z.; Driever, W. Mutations affecting development of the zebrafish retina. Development. 1996, 123, 263–273.

48. Mandyam, C.D; Harburg, G.C.; Eisch, A.J. Determination of key aspects of precursor cell proliferation, cell cycle length and kinetics in the adult mouse subgranular zone. Neuroscience. 2007, 146(1), 108–122. doi: 10.1016/j.neuroscience.2006.12.064.

49. Marcucci, F.; Murcia-Belmonte, V.; Wang, Q.; Coca, Y.; Ferreiro-Galve, S.; Kuwajima, T.; Khalid, S.; Ross, M.E.; Mason, C.; Herrera, E. The ciliary margin zone of the mammalian retina generates retinal ganglion cells. Cell Rep. 2016, 17, 3153–3164. doi: 10.1016/j.celrep.2016.11.016.

50. Marcus, R.C.; Delaney, C.L; Easter, S.S. Jr. Neurogenesis in the visual system of embryonic and adult zebrafish (Danio rerio). off. Vis. Neurosci. 1999, 16(3), 417–424. doi: 10.1017/s095252389916303x.

51. Meyer, R.L. Evidence from thymidine labeling for continuing growth of retina and tectum in juvenile goldfish. Exp. Neurol. 1978, 59, 99–111. doi: 10.1016/0014-4886(78)90204-2.

52. Miles, A.; Tropepe, V. Retinal stem cell ‘retirement plans’: growth, regulation and species adaptations in the retinal ciliary marginal zone. Int. J. Mol. Sci. 2021, 22, 6528. doi:10.3390/ijms22126528

53. Morris, A.C.; Scholz, T.L.; Brockerhoff, S.E.; Fadool, J.M. Genetic dissection reveals two separate pathways for rod and cone regeneration in the teleost retina. Dev. Neurobiol, 2008a, 68(5), 605–619. https://doi.org/10.1002/dneu.20610

54. Morris, A.C.; Scholz, T.; Fadool, J.M. Rod progenitor cells in the mature zebrafish retina. Adv. Exp. Med. Biol., 2008b, 613, 361–368. doi:10.1007/978-0-387-74904-4_42

55. Moshiri, A.; Close, J.; Reh, T.A. Retinal stem cells and regeneration. Int. J. Dev. Biol. 2004, 48, 1003–1014. doi: 10.1387/ijdb.041870am.

56. Nagashima, M.; Barthel, L.K.; Raymond, P.A. A self-renewing division of zebrafish Müller glial cells generates neuronal progenitors that require N-cadherin to regenerate retinal neurons. Development. 2013, 140, 4510–4521. doi: 10.1242/dev.090738. doi: 10.1242/dev.090738.

57. Negishi, K.; Stell, W.K.; Takasaki, Y. Early histogenesis of the teleostean retina: studies using a novel immunochemical marker, proliferating cell nuclear antigen (PCNA/cyclin). Brain Res. Dev. 1990, 55(1), 121–125.

58. Okada, T.S. Cellular metaplasia or transdifferentiation as a model for retinal cell differentiation. Curr. Top. Dev. Biol. 1980, 16, 349–380.

59. Otteson, D.C; D’Costa, A.R.; Hitchcock, P.F. Putative stem cells and the lineage of rod photoreceptors in the mature retina of the goldfish. Dev. Biol. 2001, 232(1), 62–76. doi: 10.1006/dbio.2001.0163.

60. Perron, M.; Harris, W.A. Retinal stem cells in vertebrates. Bioessays 2000, 22(8), 685–688. doi: 10.1002/1521-1878(200008)22:8<685::AID-BIES1>3.0.CO;2-C.

61. Raymond, P.A.; Rivlin, P.K. Germinal cells in the goldfish retina that produce rod photoreceptors. Dev. Biol. 1987, 122(1), 120–138. doi: 10.1016/0012-1606(87)90338-1.

62. Raymond, P.A.; Barthel, L.K., Bernardos, R.I., Perkowski, J.J. Molecular characterization of retinal stem cells and their niches in adult zebrafish. BMC Develop. Biol. 2006, 6, 36. doi: 10.1186/1471-213X-6-36.

63. Reh, T.A.; Fischer, A.J. Stem cells in the vertebrate retina. Brain Behav. Evol. 2001, 58(5), 296–305. doi: 10.1159/000057571.

64. Sánchez-Farías, N.; Candal, E. Doublecortin widely expressed in the developing and adult retina of sharks. Exp. Eye Res. 2015, 134, 90–100. doi: 10.1016/j.exer.2015.04.002.

65. Sánchez-Farías, N.; Candal, E. Identification of radial glia progenitors in the developing and adult retina of sharks. Front. Neuroanat. 2016, 10, 65. doi: 10.3389/fnana.2016.00065.

66. Stenkamp D.L. The rod photoreceptor lineage of teleost fish. Prog. Retin. Eye Res., 2011, 30(6), 395–404. doi:10.1016/j.preteyeres.2011.06.004

67. Than-Trong, E.; Bally-Cuif, L. Radial glia and neural progenitors in the adult zebrafish central nervous system. Glia. 2015, 63(8), 1406–1428. doi: 10.1002/glia.22856. doi: 10.1002/glia.22856.

68. Tropepe, V.; Coles, B.L.; Chiasson, B.J.; Horsford, D.J.; Elia, A.J.; McInnes, R.R.; van der Kooy, D. Retinal stem cells in the adult mammalian eye. Science. 2000, 287(5460), 2032–2036. doi: 10.1126/science.287.5460.2032.

69. Van Houcke, J.; Geeraerts, E.; Vanhunsel, S.; Beckers, A.; Noterdaeme, L.; Christiaens, M.; Bollaerts; I., De Groef, L.; Moons, L. Extensive growth is followed by neurodegenerative pathology in the continuously expanding adult zebrafish retina. Biogerontology. 2019, 20(1), 109–125. doi: 10.1007/s10522-018-9780-6.

70. Velasco, A.; Cid, E.; Ciudad, J.; Orfao, A.; Aijón, J.; Lara, J.M. Temperature induces variations in the retinal cell prolifer-ation rate in a cyprinid. Brain Res. 2001, 913, 190–194. doi:10.1016/S0006-8993(01)02804-9.

71. Villar-Cheda, B.; Abalo, X.M.; Villar-Cerviño, V.; Barreiro-Iglesias, A.; Anadón, R.; Rodicio, M.C. Late proliferation and photoreceptor differentiation in the transforming lamprey retina. Brain Res. 2008, 1201, 60–67. doi: 10.1016/j.brainres.2008.01.077.

72. Wan, Y.; Almeida, A.D.; Rulands, S.; Chalour, N.; Muresan, L.; Wu., Y.; Simons, B.D.; He, J.; Harris, W.A. The ciliary marginal zone of the zebrafish retina: clonal and time-lapse analysis of a continuously growing tissue. Development. 2016, 143, 1099–1107. doi: 10.1242/dev.133314.

73. Weber, I.P.; Ramos, A.P.; Strzyz, P.J.; Leung, L.C.; Young, S.; Norden, C. Mitotic position and morphology of committed precursor cells in the zebrafish retina adapt to architectural changes upon tissue maturation. Cell Rep. 2014, 7(2), 386–397. doi: 10.1016/j.celrep.2014.03.014.

74. Zerjatke, T.; Gak, I.A.; Kirova, D.; Fuhrmann, M.; Daniel, K.; Gonciarz, M.; Müller, D.; Glauche, I.; Mansfeld, J. Quanti-tative cell cycle analysis based on an endogenous all-in-one reporter for cell tracking and classification. Cell Rep. 2017, 19(9), 1953–1966. doi: 10.1016/j.celrep.2017.05.022.

75. Zupanc, G.K.H. Adult neurogenesis in the central nervous system of teleost fish: from stem cells to function and evolution. J. Exp. Biol. 2021, 224(8), jeb226357. doi: 10.1242/jeb.226357.

76. Zupanc, G.K.; Sîrbulescu, R.F. Adult neurogenesis and neuronal regeneration in the central nervous system of teleost fish. Eur. J. Neurosci. 2011, 34(6), 917–929. doi: 10.1111/j.1460-9568.2011.07854.x.

