## Supplementary material for "Decline in constitutive proliferative activity in the zebrafish retina with ageing": Table S1

| **Species** | | **Stages/ages/sizes studied** | **Maximum size/age of specimens of each species** | **Method to label proliferating or mitotic cells** | **Retinal region of analysis** | **Quantification and comparison between different life stages or ages** | | **Reference** |
| --- | --- | --- | --- | --- | --- | --- | --- | --- |
| *Haplochromis burtoni*  (African cichlid) | | Juveniles and adults | 12-15 cm | H^3^-Thymidine | ONL | No (Qualitative descriptions: “dividing cells in the ONL were easier to demonstrate in younger fish”) | | Johns and Fernald, 1981 |
|  |  | Adults (3.3 cm) | 12-15 cm | BrdU and PCNA | ONL | No | | Mack and Fernald, 1995 |
|  |  | Adults (3 cm) | 12-15 cm | PCNA | ONL | No | | Kwan et al., 1996 |
|  |  | Adults (3.5 cm) | 12-15 cm | BrdU and PCNA | ONL | No | | Mack and Fernald, 1997 |
| *Carassius auratus (goldfish)* | | Juveniles (3-6 cm) | 12-22 cm to 45 cm  30 years | Methyl- H3-Thymidine | Whole retina | No | | Meyer, 1978 |
|  |  | Juveniles and adults | 12-22 cm to 45 cm  30 years | H3-Thymidine | ONL | No (Qualitative descriptions: “dividing cells in the ONL were easier to demonstrate in younger fish”) | | Johns and Fernald, 1981 |
|  |  | Juveniles and adults (3-13 cm) | 12-22 cm to 45 cm  30 years | H3-Thymidine | ONL | No | Johns, 1982 | |
|  | Late larvae/early juveniles (20 to 51 days post-hatching) | | 12-22 cm to 45 cm  30 years | H3-Thymidine | INL and ONL | Yes (although no statistical analyses were used to compare different ages and only 1 or 2 sections were analysed per retina) | Raymond and Rivlin, 1987 | |
|  | Embryos, larvae [hatching (H0) and 1-14 days post-hatching] and adults | | 12-22 cm to 45 cm  30 years | PCNA | Whole retina | No | Negishi et al., 1990 | |
|  | Adults (2-6 cm) | | 12-22 cm to 45 cm  30 years | BrdU | ONL | No | Stenkamp et al., 1997 | |
|  | Juveniles (2.5-3.8 cm) | | 12-22 cm to 45 cm  30 years | BrdU and pH3 | Whole retina | No | Otteson et al., 2001 | |
|  | Adults (9-12 cm) | | 12-22 cm to 45 cm  30 years | PCNA | Whole retina | No | Cid et al., 2002 | |
| *Salmo trutta* (trout) | Larvae (newly hatched and 3-6 weeks post-hatching), early (3 months post-hatching) and late (1-2 years post-hatching) juveniles | | 40-80 cm to 140 cm  2-3 years (sexual maturity)  8 to 20 years | Heidenhain's haematoxylin and eosin, Ehrlich's haematoxylin and eosin, and Mallory stainings | Whole retina | No | Lyall, 1957 | |
| *Onchorynchus mykiss* (rainbow trout) | Early (2 months post-hatching) to late (2 years post-hatching) juveniles | | 3-6 mm (egg) to 12-20 mm (hatched)  2-3 years | PCNA, BrdU and IdU | INL | No (authors show that the density of PCNA+ cells in the INL decreases as eye diameter increases but without any statistical comparison) | Julian et al., 1998 | |
| *Oryzias latipes* (medaka) | Embryos, larvae [ hatching (H0) and 1-14 days post-hatching] and adults | | 3-6 cm  2 to 3-5 years | PCNA | Whole retina | No | Negishi et al., 1990 | |
| *Tinca tinca* (tench) | Adults | | 25-30 cm  20 years | PCNA | Whole retina | No | Velasco et al., 2001 | |
|  | Adults (13-16 cm) | | 25-30 cm  20 years | PCNA | Whole retina | No | Cid et al., 2002 | |
| *Danio rerio* (zebrafish) | Embryos (24 and 48 hpf) and adults (6-8 months postfertilization) | | Sexual maturation (3 months)  2/3 to 5 years old | BrdU | Whole retina | No (qualitative descriptions: “the number of labelled cells is greater in the embryos”) | Marcus et al., 1999 | |
|  | Adults (3.5-4 cm) | | Sexual maturation (3 months)  2/3 to 5 years old | PCNA | Whole retina | No | Cid et al., 2002 | |
|  | Juveniles (1 to 2 mpf) | | Sexual maturation (3 months)  2/3 to 5 years old | BrdU and PCNA | Central retina | No | Bernardos et al., 2007 | |
|  | Adults (6 to 48 mpf) | | Sexual maturation (3 months)  2/3 to 5 years old | PCNA | CMZ | Yes (decreased proliferation with ageing) | Van Houcke et al., 2019 | |
